# Periodic and aperiodic changes to cortical EEG in response to pharmacological manipulation

**DOI:** 10.1101/2023.09.21.558828

**Authors:** Sofia V. Salvatore, Peter M. Lambert, Ann Benz, Nicholas R. Rensing, Michael Wong, Charles F. Zorumski, Steven Mennerick

**Author notes:** Correspondence to: Steven Mennerick, Department of Psychiatry, Washington University in St. Louis School of Medicine, 660 S. Euclid Ave., MSC 8134-0181-0G, St. Louis, MO 63110.

## Abstract

Cortical electroencephalograms (EEG) may help understanding of neuropsychiatric illness and new treatment mechanisms. The aperiodic component (1/*f*) of EEG power spectra is often treated as noise, but recent studies suggest that changes to the aperiodic exponent of power spectra may reflect changes in excitation/inhibition (E/I) balance, a concept linked to antidepressant effects, epilepsy, autism, and other clinical conditions. One confound of previous studies is behavioral state, because factors associated with behavioral state other than E/I ratio may alter EEG parameters. Thus, to test the robustness of the aperiodic exponent as a predictor of E/I ratio, we analyzed active exploration in mice using video EEG following various pharmacological manipulations with the Fitting Oscillations & One Over F (FOOOF) algorithm. We found that GABA_A_ receptor (GABA_A_R) positive allosteric modulators increased the aperiodic exponent, consistent with the hypothesis that an increased exponent signals enhanced cortical inhibition, but other drugs (ketamine and GABA_A_R antagonists at sub-convulsive doses) did not follow the prediction. To tilt E/I ratio more selectively toward excitation, we suppressed the activity of parvalbumin (PV) interneurons with Designer Receptors Exclusively Activated by Designer Drugs (DREADDs). Contrary to our expectations and studies demonstrating increased cortical activity following PV suppression, circuit disinhibition with the DREADD increased the aperiodic exponent. We conclude that the aperiodic exponent of EEG power spectra does not yield a universally reliable marker of E/I ratio. Alternatively, the concept of E/I state may be sufficiently oversimplified that it cannot be mapped readily onto an EEG parameter.

**Significance StateBment:** Neuropsychiatric illness is widely prevalent and debilitating. Causes are not well understood, but some hypotheses point toward altered excitation/inhibition (E/I) balance. Here, we use cortical electroencephalograms (EEG) in mice, given applicability of cortical EEG across species, and evaluate the impact of validated drugs, including anxiolytics (pentobarbital and diazepam), along with novel rapid-acting antidepressants (ketamine and allopregnanolone). We focus on analyzing the aperiodic component of EEG power spectra, which may be associated with changes in E/I ratio. We show that aperiodic exponent of EEG power spectra is not a reliable marker of E/I ratio. Moreover, the concept of E/I ratio may be too broad and complex to be defined by an EEG parameter.

## Introduction

Analysis of cortical electroencephalograms (EEG) may help drive understanding of neuropsychiatric illness and of new treatment mechanisms. Neuronal oscillations in EEG recordings characterize various behavioral states, including perception, cognition, social interaction, and neuropsychiatric disorders (Buzsáki and Draguhn, 2004; Kehrer, 2008; Sgadò et al., 2011; Voytek and Knight, 2015; Fitzgerald and Watson, 2018). The aperiodic component (1/*f*) of EEG power spectra is classically treated as noise but has come under scrutiny recently for its potential significance in disease (Donoghue et al., 2020). Specifically, changes to the exponent of the aperiodic component of power spectra are associated with excitation/inhibition (E/I) ratio, a concept linked to antidepressant effects of new treatments (disinhibition), to epilepsy (over-excitation), and to developmental disorders such as autism (Miller et al., 2016; Bozzi et al., 2018; Ahmad et al., 2022). Here we seek to test the validity and utility of the aperiodic analysis using drugs with known underlying mechanisms of action.

GABA_A_ receptor positive allosteric modulators (PAMs), such as pentobarbital and diazepam, are benchmark compounds that globally increase inhibition. GABA_A_R antagonists such as bicuculine and picrotoxin have long been used to model seizures and epilepsy and thus may serve as benchmark drugs for excitatory effects (Schechter and Tranier, 1977; Freund et al., 1987; Singh et al., 2010; Johnston, 2013). Effects on sub-convulsive doses of picrotoxin and bicuculline *in vivo* are unknown but we hypothesize increase E/I ratio.

Ketamine represents a newer generation of antidepressant treatments. It is an antagonist of NMDA-type glutamate receptors (Zarate et al., 2006) that initiates durable antidepressant effects at low doses and is a dissociative anesthetic at high doses (Miller et al., 2016). Low doses of ketamine may cause cortical disinhibition (Monyer et al., 1994; Moghaddam et al., 1997; Seamans, 2008; Zanos and Gould, 2018). Recently, ketamine was found to decrease the aperiodic exponent in humans at anesthetic doses, which could be consistent with the disinhibition hypothesis (Waschke et al., 2021). However, whether this result pertains to low doses where behavioral state can be held constant remains unknown.

Another durable, rapid antidepressant is allopregnanolone (AlloP), an endogenous neurosteroid and potent GABA_A_ receptor (GABA_A_R) PAM. Like ketamine, disinhibition may explain AlloP’s antidepressant effects (Lambert et al., 2022) through subunit-selective GABA_A_R effects on interneurons (Antonoudiou et al., 2022). AlloP effects on the aperiodic component of cortical EEG are unknown. We previously found that AlloP’s effects on the periodic component of cortical EEG signal was indistinguishable from pentobarbital but distinct from diazepam (Lambert et al., 2023). Here we test whether AlloP’s impact on non-oscillatory EEG structure may be distinct from the other two GABA_A_R PAMs and consistent with disinhibition.

Past studies of pharmacological effects on the aperiodic component of EEG have focused on anesthesia (Gao et al., 2017; Waschke et al., 2021), where behavioral state represents a confounding variable. Using video-EEG recordings in freely behaving mice, we focus on active exploration, and we canvass the above drugs as well as cell-type selective manipulation of interneurons with Designer Receptors Exclusively Activated by Designer Drugs (DREADDs). We assess each manipulation against the hypothesis that the exponent of the aperiodic fit reveals cortical E/I ratio in the normal mouse brain; an increase in exponent is indicative of stronger cortical inhibition and vice-versa for a decrease in exponent. We analyze our neural data using a recently developed algorithm, fitting oscillations & one over f (FOOOF) (Donoghue et al., 2020). Overall, although several pharmacological agents (notably GABA_A_R PAMs) were consistent with the hypothesis that a steeper (larger) aperiodic exponent depicts inhibition, other drugs did not follow the prediction. We conclude that analysis of the aperiodic exponent of EEG power spectra does not yield a reliable marker of E/I state. Alternatively, the concept of a circuit E/I state may be sufficiently oversimplified that it cannot be mapped cleanly onto an EEG parameter.

## Materials and Methods

### Ethical approval

All procedures were carried out in accordance with National Institute of Health (NIH) guidelines and approved by the Washington University Institutional Animal Care and Use Committee, protocol 22-0344. Pain and suffering were alleviated with appropriate anesthesia and analgesia during surgical procedures. Animals were obtained from the Jackson Laboratory or born in our colony and were reared under the care of the Washington University School of Medicine Division of Comparative Medicine. Animals had ad libitum access to food and water. Mice were euthanized at the end of studies according to NIH guidelines for minimizing pain.

### EEG surgery

7-9 week old mice (from a C57BL6/J JAX# 000664 background) of both sexes were surgically implanted with electrodes and EMG as previously described (Lambert et al., 2023). For our behavioral state experiment, we followed the same procedure described above, but we used *Gabrd* floxed mice from a mixed C57BL6/J and 129S1/SvImJ background (JAX #023836), due to colony accessibility.

### EEG recording

Mice were maintained on a reverse lighting cycle and recordings were initiated in the first half of the dark cycle to enrich for active wake behaviors during the period of acute drug exposure. EEG was acquired as previously published (Lambert et al., 2023). A series of 12 cohorts of 4 animals each were recorded for a total of 4-15 animals per drug group (pentobarbital 5M/2F, diazepam 3M/2F, picrotoxin 2M/2F, bicuculline 5M/1F, sub-anesthetic ketamine 7M/8F, anesthetic ketamine 2M/2F, AlloP 3M/2F, DCZ hM4D(Gi) 3M/2F, DCZ hM3D(Gq) 1M, 5F). For behavioral state experiments, 1 cohort of 4 animals was recorded (3M/1F). Recordings for the experimental session began with a 30-minute baseline recording period. Next, a vehicle injection for each drug condition was delivered, followed 30 minutes later by the active drug. EEG monitoring continued for 6 hours. For DREADD experiments, recordings began with a 45-minute baseline period, followed by the vehicle or active drug. Animals in DREADD experiments were recorded for 2 sessions, one with vehicle and one with active drug (order randomized). Baseline EEG was not affected by DREADD activation after a 48-h washout. Experiments that canvassed animals sleep as well as active wake states recorded EEG for 12 h in the light phase. All recordings were accompanied by video film, captured using a monochrome camera with an infrared lens and Bonsai software.

### Drugs

Pentobarbital (Sigma) and ketamine (Sigma) were both dissolved in sterile saline (0.9% NaCl) to final concentrations of 6 mg/mL and 2 mg/mL, respectively. Diazepam (Sigma) was dissolved in 40% propylene glycol in sterile saline at a concentration of 0.2 mg/mL. Picrotoxin (Tocris) was initially dissolved in 45% 2-hydroxypropyl β-cyclodextrin (CDX) at a concentration of 1 mg/mL and sonicated until completely dissolved, then further diluted in sterile saline to 0.6 mg/mL and 22.5% CDX. (+)-Bicuculline (Tocris) was dissolved in 10% 0.1 N HCl and sterile saline at a concentration of 0.1 mg/mL. AlloP (Sigma) was initially dissolved in 45% CDX at a concentration of 1.2 mg/mL and sonicated until completely dissolved, then further diluted in sterile saline to 0.6 mg/mL AlloP and 22.5% CDX. Deschloroclozapine (DCZ) (Tocris) was dissolved in 0.5% DMSO in sterile saline at a concentration of 0.01 mg/mL. All drugs were delivered as a single intraperitoneal injection with the following doses (mg/kg): pentobarbital 15, diazepam 1, picrotoxin 2, bicuculline 1, ketamine 10 and 120, AlloP 5, DCZ 0.1. Vehicle injections were of the same solutions without the drug, for each treatment. These doses were determined by pilot studies of escalating doses to be subanesthetic or subconvulsive as appopriate.

### DREADDs

4-6 week old parvalbumin (PV)-cre heterozygous mice from a C57BL6/J background (JAX #017320) were injected retro-orbitally with pAAV-hSyn-DIO-hM4D(Gi)-mCherry (Addgene, 44362-PHPeB) (Gi) or pAAV-hSyn-DIO-hM3D(Gq)-mCherry (Gq) (Addgene, 44361-PHPeB)(Chan et al., 2017) and incubated for at least 4 weeks prior to injection with DCZ. DCZ was chosen as the agonist for DREADDs due to its potent, selective, metabolically stable, and fast-acting effects that are better than the classically used clozapine-*N-*oxide (CNO) (Nagai et al., 2020). Pilot studies revealed that a dose of 0.03 mg/kg DCZ produced negligible EEG effects on hM4D(Gi) mice. Thus, we selected a higher dose of 0.1 mg/kg, which has also shown strong effects on hM3D(Gq) expressed in GABAergic neurons (Ferrari et al., 2022).

### Immunostaining

Mice were perfused transcardially with PBS for 3 min, followed by 4% paraformaldehyde for 4 min. Brains were post-fixed in 4% paraformaldehyde overnight at 4LC and cryoprotected for 48h in 30% sucrose at 4LC. Brains were frozen on dry ice, then sagittally sectioned at 45 μm on a freezing microtome. Free-floating sections were blocked in 3% normal goat serum, 0.3% Triton-X detergent, and 2% BSA in PBS for 1 h at room temperature. Then they were incubated in primary rat anti-mCherry antibody (Invitrogen, M11217) and primary rabbit anti-PV (Swant, PV28) diluted 1:1000 in block solution at 4LC, overnight and shaking. Subsequently, sections were washed 3 times with PBS and incubated in secondary Alexa Fluor 555 conjugated goat anti-rat (Invitrogen, A-21434) and secondary Alexa Fluor 488 conjugated goat anti-rabbit (Invitrogen, A-11008) diluted 1:500 in PBS for 2 h at room temperature, in the dark with shaking. Finally, sections were washed 3 times with PBS and mounted on slides for imaging.

### Microscopy

Images for quantification of viral expression were taken on a Nikon Eclipse TE2000-S microscope equipped with epi-fluorescence illumination and a camera (Photometrics, CoolSNAP ES^2^). Images were acquired with Micro-Manager software. 10x images from 3 different fields were taken for each mouse. Image capture order: phase, Alexa Fluor 488, mCherry. %mCherry/PV fluorescence was quantified using the analyze particles tool in FIJI. For each mouse, 3 fields were quantified and averaged together.

Widefield imaging was performed on a Zeiss AxioScan Z1 equipped with a mercury illumination source and a Hammatsu Orca Flash sCMOS camera, at 20x.

### EEG analysis

Raw data were imported into MATLAB for further analysis. Time frequency spectrograms were generated from a wavelet transform of the raw EEG signal, as previously described (Lambert et al., 2023).

Power spectra were created for each condition (baseline, vehicle, drug) using the 5 minutes of active wake from the parietal right electrode. EEG was analyzed using the segmented continuous multi-taper FFT method from the Chronux toolbox in MATLAB. EEG data was segmented into 5 second windows and multi-taper parameters were [TW, K] = 7.5, 14. Power spectra were binned to a spectral resolution of 0.6 Hz before plotting.

For experiments that assessed animals’ baseline power spectra across behavioral states, EEG/EMG was scored into active wake, REM and NREM stages. This was done in a semi-automated manner, using the AccuSleep toolbox in MATLAB, as previously described (Lambert et al., 2023). Having identified EEG from these diverse states, 10 minutes of each state were used to create power spectra, as detailed above.

Power spectra were parameterized into periodic and aperiodic components using FOOOF in Python. Across all conditions, the FOOOF parameters were: peak_width_limits = [2,40], aperiodic_mode = ‘knee’, and the frequency reported included 2-100 Hz. Knee frequency corresponds to the inflection frequency on log-log plots of the aperiodic component (Donoghue et al., 2020), representing the idea that there is not a constant 1/*f* relationship across the entire range of frequencies. We excluded 1Hz due to its impediment in creating accurate aperiodic fits – particularly in the high frequency range. Across drugs, the min_peak_height and max_n_peaks parameters were altered to best fit the power spectra of each drug. These parameters were maintained within drugs and across all behavioral states.

Raw parietal EEG power spectra of drugs previously published (pentobarbital, diazepam, ketamine sub-anesthetic N = 5, and AlloP) can be found in (Lambert et al., 2022b; Supplementary Fig. 1). Raw parietal EEG power spectra of new treatments (picrotoxin, behavioral states, sub-anesthetic ketamine with addition of 10 animals, anesthetic ketamine, DCZ in DREADD expressing animals) can be found in Extended Data Figure 3-1.

### Statistical analysis and figure preparation

EEGs from animals treated with pentobarbital, diazepam, AlloP and those of 5 animals treated with ketamine were previously recorded and published using a different analysis that focused on the periodic component (Lambert et al., 2023). Here, we focus on FOOOF to analyze the aperiodic component of the power spectra from these data. We further expand the analysis to different pharmacological agents, including anesthetic ketamine, picrotoxin, and DREADDs. If a theta or gamma oscillation fell below detection limit of FOOOF in any treatment, that animal was excluded from that frequency band’s statistical analysis tests. Some knee frequencies could not be calculated due to small (>-1) negative knee values, and they are evident in figures as missing points on knee graphs (e.g., ketamine data).

For the ketamine dataset, initial analysis of the 5 mice previously recorded showed a trend toward decreased aperiodic exponent (repeated measures one-way ANOVA, F (1.293, 5.172) = 1.566 p = 0.2771). Given the large effect size between vehicle and ketamine of 1.01, we conducted a two-tailed t-test power analysis to determine the minimum required sample size to achieve 95% of power, with an α = .05. The minimum sample size needed for this effect size was N = 15, (t = 2.14479, df = 14). Thus, we injected 10 additional mice for a total N of 15 mice analyzed herein.

GraphPad Prism was used to plot graphs and perform statistical analyses. Error bars represent mean ± standard error of mean (SEM). For presentation of images, maximum and minimum gray values were adjusted in FIJI and assembled in InkScape.

## Results

### GABA_A_R PAMs increase aperiodic exponent as expected

We focused on EEG changes during active (exploratory) wake for baseline, vehicle, and acute drug action (∼1h post injection). We used data previously analyzed by different methods (Lambert et al., 2022b; Figure 2) to evaluate the impact of a classic GABA_A_R PAM (pentobarbital) on parietal EEG periodic and aperiodic components. The characteristic oscillatory features in power spectra of active wake EEG, including prominent theta frequency peak and a broad gamma frequency peak, were adequately described by the FOOOF algorithm (Fig. 1A baseline and vehicle). Similarly, the impact of a single intraperitoneal injection of 15 mg/kg of pentobarbital on overall EEG structure was well fit (Figs. 1A, PTB condition). Later figures have comparably good fits unless otherwise indicated. Subtracting oscillatory components revealed the aperiodic fit (Fig.1A, dashed lines). Extracting the oscillatory components from the fits revealed changes to the theta, beta, and gamma oscillatory components (Fig. 1B) similar to those reported previously by other methods (Lambert et al., 2023). Importantly, the aperiodic component of the drug condition exhibited a steeper slope (larger exponent) beyond the inflection (knee) frequency (Donoghue et al., 2020)(Fig. 1C). This shift is consistent with the hypothesis that reducing E/I ratio increases the aperiodic exponent. Pentobarbital also surprisingly increased the offset (y-axis intercept) of the aperiodic fit, which is correlated with neuronal population spiking (Donoghue et al., 2020)(Fig.1C).

**Figure 1.**
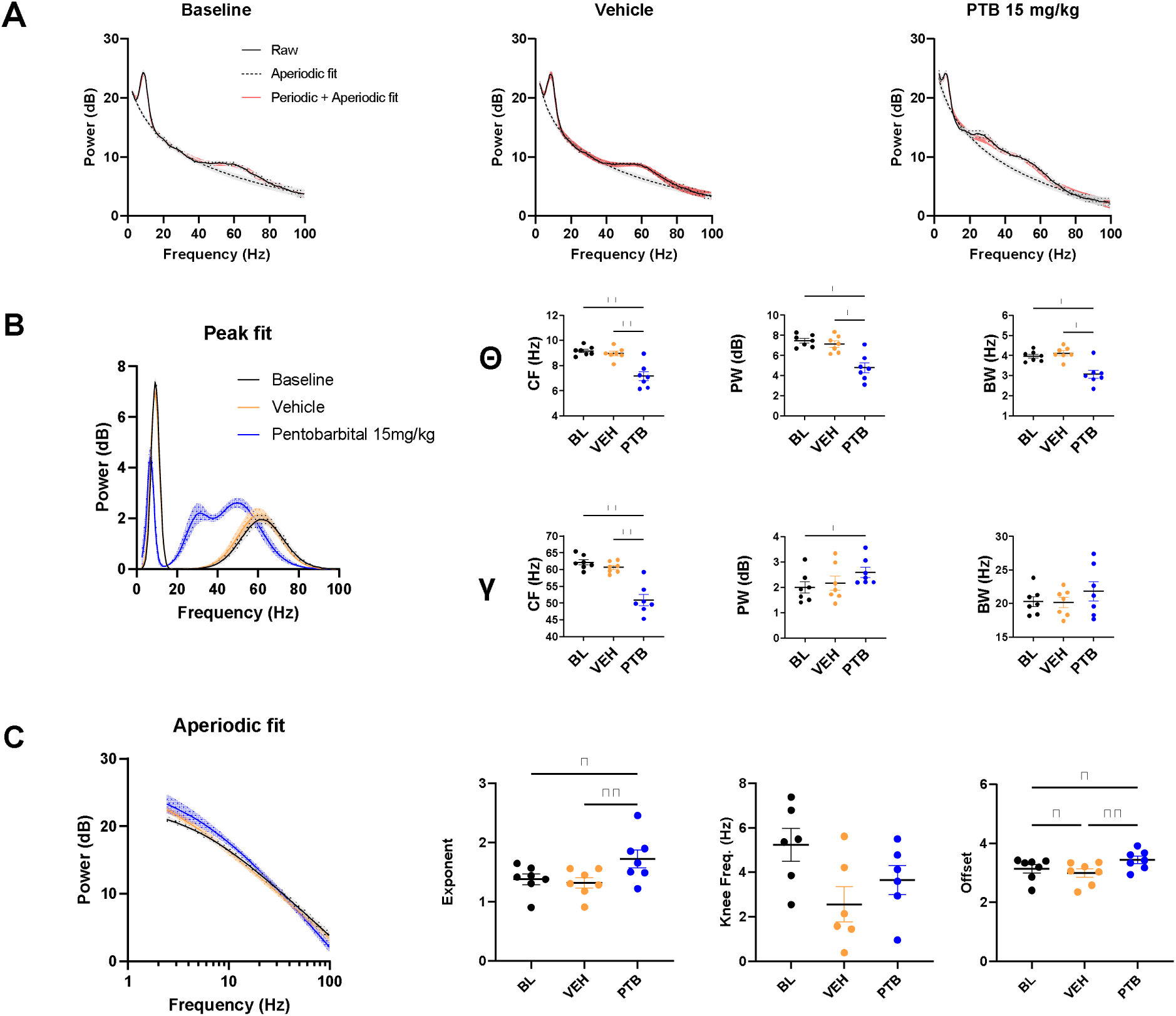
Pentobarbital 15 mg/kg, a broad spectrum GABA_A_R PAM, increases network inhibition. **(A)** Raw power spectra of baseline (BL), vehicle (VEH), and pentobarbital (PTB), accurately fit by FOOOF: aperiodic and periodic+aperiodic components. **(B)** Periodic fit of peaks from baseline, vehicle, and PTB effects on parietal EEG raw power spectra. Emergence of a beta (12-30) peak with PTB. Parameters of the theta (1-5) and gamma (30-100) peaks: peak central frequency (CF), peak power (PW) and peak bandwidth (BW). For the theta peak, a repeated measures one-way ANOVA revealed an effect of treatment on CF (F (1.08, 6.481) = 22.14 p = 0.0025). Tukey’s multiple comparison showed a difference between BL and PTB CF (**p = 0.0064) and VEH and PTB CF (**p = 0.0090). Repeated measures one-way ANOVA revealed an effect of treatment on PW (F (1.104, 6.626) = 15.12 p = 0.0061). Tukey’s multiple comparison showed a difference between BL and PTB PW (*p = 0.0.151) and VEH and PTB PW (*p = 0.0217). Repeated measures one-way ANOVA revealed an effect of treatment on BW (F (1.081, 6.487) = 16.51 p =0.0052). Tukey’s multiple comparison showed a difference between BL and PTB BW (*p = 0.0142) and VEH and PTB BW (*p = 0.0150). For the gamma peak, a repeated measures one-way ANOVA revealed an effect of treatment on CF (F (1.162, 6.969) = 25.41p = 0.0012). Tukey’s multiple comparison showed a difference between BL and PTB CF (**p = 0.0046) and VEH and PTB CF (**p = 0.0057). Repeated measures one-way ANOVA revealed an effect of treatment on PW (F (1.050, 6.301) = 6.588 p = 0.0398). Tukey’s multiple comparison showed a difference between BL and PTB PW (*p = 0.0127). Repeated measures one-way ANOVA revealed no effect of treatment on BW (F (1.249, 7.493) = 1.412 p =0.2825). **(C)** Aperiodic fit of baseline, vehicle, and pentobarbital effects on parietal active wake EEG raw power spectra. Parameters of the aperiodic fit: exponent, knee frequency (knee freq.), and offset. Repeated measures-one way ANOVA showed an effect of treatment on exponent (F (1.100, 6.597) = 16.25 p =0.0051). Tukey’s multiple comparison showed a difference between BL and PTB exponent (*p = 0.0288) and VEH and PTB exponent (**p = 0.0076). Repeated measures one-way ANOVA revealed an effect of treatment on offset (F (1.437, 8.620) = 24.52 p = 0.0005). Tukey’s multiple comparison showed a difference between BL and VEH offset (*p = 0.0225), BL and PTB offset (*p = 0.0166), and VEH and PTB offset (**p= 0.0023).

**Figure 2.**
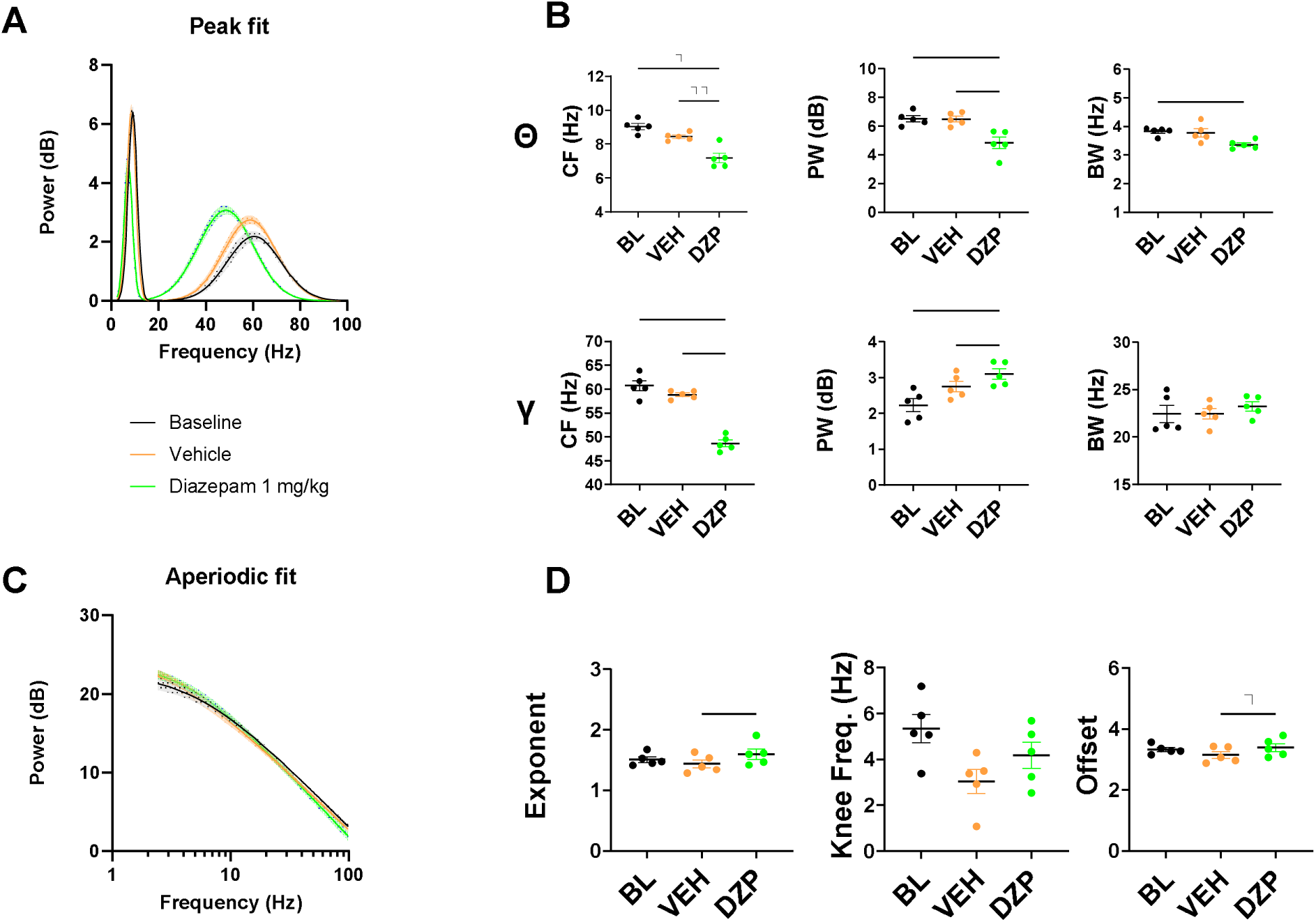
Validation of FOOOF with another network inhibitor, diazepam 1 mg/kg. **(A)** Periodic fit of peaks from baseline (BL), vehicle (VEH), and diazepam (DZP) effects on parietal active wake EEG raw power spectra. **(B)** Parameters of theta (1-5) and gamma (30-100) peaks: peak central frequency (CF), peak power (PW) and peak bandwidth (BW). For the theta peak, a repeated measures one-way ANOVA revealed an effect of treatment on CF (F (1.036, 4.144) = 18.36 p = 0.0117). Tukey’s multiple comparison showed a difference between BL and DZP CF (*p = 0.0286) and VEH and DZP CF (**p = 0.0088). Repeated measures one-way ANOVA revealed an effect of treatment on PW (F (1.912, 7.646) = 23.05 p = 0.0006). Tukey’s multiple comparison showed a difference between BL and DZP PW (**p = 0.0074) and VEH and DZP PW (*p = 0.0133). Repeated measures one-way ANOVA revealed a trend towards an effect of treatment on BW (F (1.484, 5.935) = 5.488 p = 0.0504). Tukey’s multiple comparison showed a difference between BL and DZP BW (*p = 0.0268). For the gamma peak, a repeated measures one-way ANOVA revealed an effect of treatment on CF (F (1.061, 4.245) = 53.92 p = 0.0014). Tukey’s multiple comparison showed a difference between BL and DZP CF (**p = 0.0050) and VEH and DZP CF (***p = 0.0004). Repeated measures one-way ANOVA revealed an effect of treatment on PW (F (1.055, 4.221) = 14.29 p = 0.0172). Tukey’s multiple comparison showed a difference between BL and DZP PW (*p = 0.0302) and VEH and DZP PW (**p = 0.0030). Repeated measures one-way ANOVA revealed no effect of treatment on BW (F (1.879, 7.518) = 0.7948 p = 0.4789). **(C)** Aperiodic fit of baseline, vehicle, and diazepam effects on parietal active wake EEG raw power spectra. **(D)** Parameters of aperiodic fit: exponent, knee frequency (knee freq.), and offset. Repeated measures one-way ANOVA revealed an effect of treatment on exponent (F (1.552, 6.207) = 7.425 p = 0.0261). Tukey’s multiple comparison showed a difference between VEH and DZP exponent (*p = 0.0358). Repeated measures one-way ANOVA revealed an effect of treatment on offset (F (1.475, 5.901) = 5.913 p = 0.0444). Tukey’s multiple comparison showed a difference between VEH and DZP offset (*p = 0.0208).

To validate the change to the aperiodic exponent with another GABA_A_R PAM, we fit power spectra from mice injected with diazepam (1 mg/kg). As previously documented with the same dataset (Lambert et al., 2022b; Figure 2), diazepam slowed the theta peak central frequency, reduced its power, and narrowed its bandwidth (Fig. 2 A-B). The central frequency of the gamma peak slowed while power increased, with no change in bandwidth (Fig. 2 A-B). For the aperiodic fit, diazepam increased the exponent and offset (Fig 2 C-D), qualitative similar to pentobarbital. Overall, GABA_A_ receptor PAMs increased the exponent of aperiodic fits as predicted by the hypothesis.

### GABA_A_R antagonists at sub-convulsive doses create unexpected sedation and fail to decrease the aperiodic exponent

Next, we tested the aperiodic exponent hypothesis with changes expected to decrease inhibition and lower the aperiodic exponent. First, we injected mice with 2 mg/kg picrotoxin, which we determined from pilot experiments to be just sub-convulsant. Picrotoxin produced an unexpected sedation of the mice (Extended Data Fig. 3-2). To account for behavioral state differences on EEG we analyzed the baseline, vehicle, sedated state (up to 45 min post injection), active wake during picrotoxin recovery (30 min – 1 h post-injection), and a wash out (∼2 h after drug). During active wake following picrotoxin, theta central frequency slowed and power decreased; theta power also decreased during sedation with no change in bandwidth (Fig. 3 A-B). For the gamma peak during active wake, picrotoxin slowed central frequency (Fig. 3 A-B). Picrotoxin produced no changes in gamma power or bandwidth (Fig. 3 A-B). Picrotoxin sedation revealed a clear increase in exponent for the aperiodic component (Fig. 3C-D). Although this might be expected for sedation, the increased exponent persisted with active wake during recovery relative to full wash out (Fig. 3 C-D). Sedation by picrotoxin also increased the aperiodic offset (Fig. 3D). To test whether picrotoxin’s sedative effects were unique to this GABA_A_R antagonist, we injected animals with bicuculline at a sub-convulsive dose of 1 mg/kg, (Freund et al., 1987; Yajima et al., 2000). Like picrotoxin, bicuculline produced immediate sedation in the 5 male mice and 1 female mouse tested. Given the confound of behavioral change, we did not perform EEG analysis.

**Figure 3.**
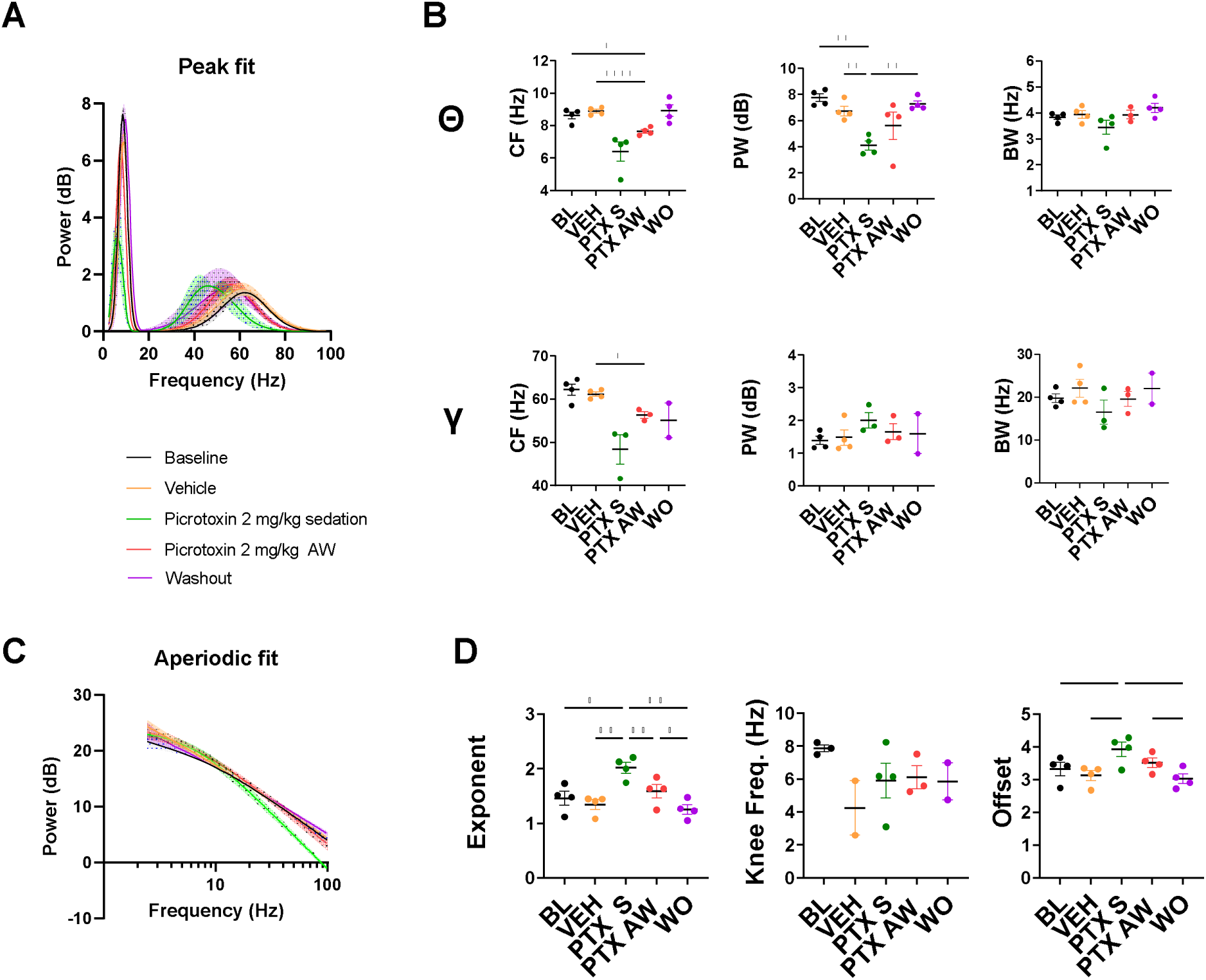
Picrotoxin at sub-convulsive dose of 2 mg/kg causes unexpected sedation; no decrease of exponent upon wake. **(A)** Periodic fit of peaks from baseline (BL), vehicle (VEH), and picrotoxin (PTX) sedation (S), PTX active wake (AW), and washout (WO) effects on parietal active wake EEG raw power spectra. **(B)** Parameters of theta (1-5) and gamma (30-100) peaks: peak central frequency (CF), peak power (PW) and peak bandwidth (BW). For the theta peak, a repeated measures one-way ANOVA revealed an effect of treatment on CF (F (1.816, 5.448) = 11.71 p = 0.0113). Tukey’s multiple comparison showed a difference between BL and PTX AW CF (*p = 0.0219) and VEH and PTX AW CF (****p =< 0.0001). Repeated measures one-way ANOVA revealed an effect of treatment on PW (F (1.268, 3.804) = 11.56 p = 0.0275). Tukey’s multiple comparison showed a difference between BL and PTX S PW (**p = 0.0073), VEH and PTX S (**p = 0.0031), and PTX S and WO (**p = 0.0035). Repeated measures one-way ANOVA revealed no effect of treatment on BW (F (1.799, 5.396) = 2.636 p = 0.1600). For the gamma peak, a mixed effects analysis revealed a trend towards an effect of treatment on CF (F (1.305, 2.611) = 8.959 p = 0.0687). Tukey’s multiple comparison showed a difference between VEH and PTX AW CF (*p = 0.0117). A mixed effects analysis revealed no effect of treatment on PW (F (0.6311, 1.262) = 1.252 p = 0.3747). A mixed effects analysis revealed no effect of treatment on BW (F (0.7340, 1.468) = 0.2835). **(C)** Aperiodic fit of baseline, vehicle, and picrotoxin sedation, picrotoxin active wake, and washout effects on parietal EEG raw power spectra. **(D)** Parameters of aperiodic fit: exponent, knee frequency (knee freq.), and offset. Repeated measures one-way ANOVA revealed an effect of treatment on exponent (F (1.724, 5.172) = 42.95 p = 0.0007). Tukey’s multiple comparison showed a difference between BL and PTX S exponent (*p = 0.0132), VEH and PTX S (**p = 0.0027), PTX S and PTX AW (**p = 0.0059), PTX S and WO (**p = 0.0018), and PTX AW and WO (*p = 0.0223). Repeated measures one-way ANOVA revealed an effect of treatment on offset (F (1.486, 4.457) = 20.23 p = 0.0068). Tukey’s multiple comparison showed a difference between BL and PTX S offset (*p = 0.0168), VEH and PTX S (*p = 0.0155), PTX S and WO (*p = 0.0280), and PTX AW and WO (**p = 0.0020).

Because of unexpected sedation with GABA_A_R antagonists, we examined whether behavioral sleep states akin to sedation/hypnosis alter the aperiodic exponent of EEG. We analyzed EEG with the FOOOF algorithm from naïve animals during a 12 h light period, categorizing states into active wake, rapid-eye movement (REM) sleep, and non-REM (NREM) sleep. For theta peak fit, REM increased overall power in oscillatory components (Extended Data Fig. 3-3 A-B). For the gamma peak, REM accelerated central frequency while NREM decreased power and decreased bandwidth (Extended Data Fig. 3-3 A-B). For the aperiodic fit, both sleep states increased the aperiodic exponent, with NREM associated with the larger effect size. The same trends followed for offset (Extended Data Fig. 3-3 C-D). Overall, because sleep influenced aperiodic components, variables associated with behavioral state should be considered when interpreting effects on aperiodic components of EEG. Sleep states are generally consistent with the hypothesis that the aperiodic exponent increases with inhibition, assuming sleep represents a state of reduced E/I ratio (Bridi et al., 2020).

### Ketamine, a putative agent of cortical disinhibition, fails to change the aperiodic exponent

We next examined the effects of sub-anesthetic ketamine (10 mg/kg) on power spectra with FOOOF, hypothesizing this sub-anesthetic dose associated with antidepressant effects would decrease the exponent according to the cortical disinhibition hypothesis, whereby ketamine selectively suppresses firing of inhibitory interneurons (Kavalali and Monteggia, 2012; Zanos and Gould, 2018). Based on a power analysis of an earlier dataset of 5 mice, (Lambert et al., 2022b; Figure 3), we acquired data from a full set of 15 mice (Figure 4) with a single intraperitoneal injection of 10 mg/kg ketamine. Ketamine slowed theta peak central frequency and widened bandwidth, with no change in power (Fig. 4 A-B). Ketamine also increased power in the gamma peak, with no effects on central frequency or bandwidth (Fig. 4 A-B). These changes in oscillatory components are all consistent with our earlier analysis (Lambert et al., 2023). However, in contrast to our expectations, there was no effect of ketamine on either the exponent or offset of the aperiodic component (Fig. 4C-D).

**Figure 4.**
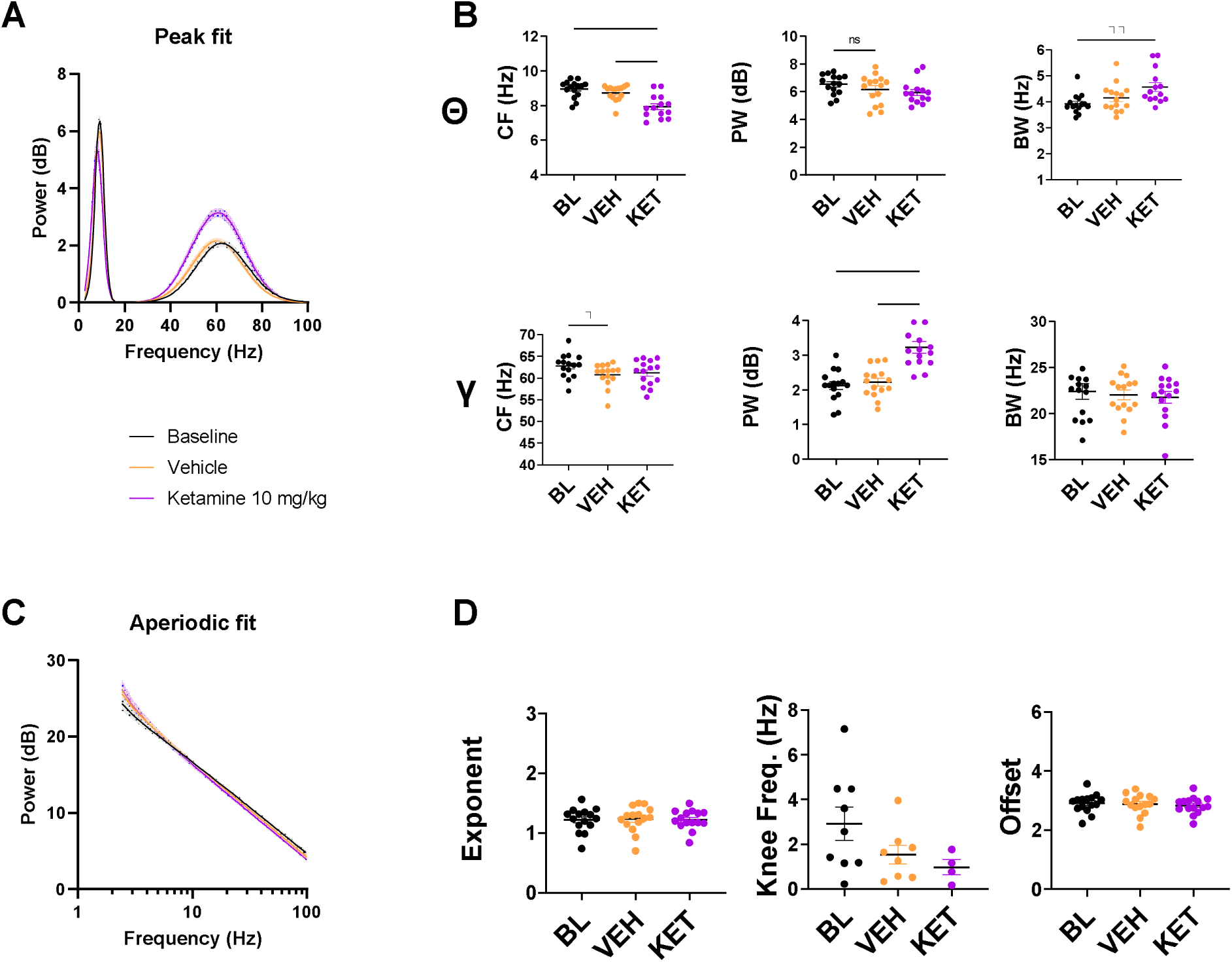
Ketamine 10 mg/kg does not alter aperiodic exponent. **(A)** Periodic fit of peaks from baseline (BL), vehicle (VEH), and ketamine (KET) effects on parietal active wake EEG raw power spectra. **(B)** Parameters of theta (1-5) and gamma (30-100) peaks: peak central frequency (CF), peak power (PW) and peak bandwidth (BW). For the theta peak, a repeated measures one-way ANOVA revealed an effect of treatment on CF (F (1.089, 15.24) = 6.251 p = 0.0223). Tukey’s multiple comparison showed a difference between BL and KET CF (*p = 0.0454). Repeated measures one-way ANOVA revealed an effect of treatment on PW (F (1.224, 17.14) = 4.546 p = 0.0412). Tukey’s multiple comparison showed a difference between BL and KET PW (*p = 0.0362). Repeated measures one-way ANOVA revealed no effect of treatment on BW (F (1.336, 18.70) = 0.6114 p = 0.4896). For the gamma peak, a repeated measures one-way ANOVA revealed no effect of treatment on CF (F (1.475, 20.64) = 2.868 p = 0.0924). Tukey’s multiple comparison showed a difference between BL and VEH CF (*p = 0.0261). Repeated measures one-way ANOVA revealed an effect of treatment on PW (F (1.503, 21.04) = 29.20 p < 0.0001). Tukey’s multiple comparison showed a difference between BL and KET PW (****p < 0.0001) and VEH and KET PW (***p = 0.0003). Repeated measures one-way ANOVA revealed no effect of treatment on BW (F (1.698, 23.77) = 0.3980 p = 0.6427). **(C)** Aperiodic fit of baseline, vehicle, and ketamine effects on parietal active wake EEG raw power spectra. **(D)** Parameters of aperiodic fit: exponent, knee frequency (knee freq.), and offset. Repeated measures one-way ANOVA revealed no effect of treatment on exponent (F (1.785, 24.99) = 0.08815 p = 0.8968). Repeated measures one-way ANOVA revealed no effect of treatment on offset (F (1.602, 22.43) = 0.8705 p = 0.4102).

Anesthetic doses of ketamine in humans decrease the exponent of the aperiodic component of EEG compared to waking rest (Waschke et al., 2021). To test these results in mice, we injected a small number of mice with an anesthetic dose of ketamine (120 mg/kg) (Irifune et al., 1992). Anesthetic ketamine produced changes in oscillatory components generally consistent with sub-anesthetic doses (Extended Data Fig. 4-1 A-B). In this limited sample, anesthetic ketamine did not statistically change the exponent or offset (Extended Data Fig. 4-1 C-D). Given the behavioral state confound, we did not pursue a larger sample size, although the trend for the aperiodic exponent was toward a decrease, consistent with the human data.

### AlloP increases aperiodic exponent, like other GABA PAMs

Another rapid-acting antidepressant compound is AlloP (brexanolone/Zulresso). We hypothesized that AlloP may decrease the aperiodic exponent as a result of disinhibition (Antonoudiou et al., 2022), distinct from other GABA_A_ PAMs. For oscillatory components, AlloP (5 mg/kg) closely resembled pentobarbital, as we observed previously with the same dataset (Lambert et al., 2022b; Figure 1, 2). AlloP slowed the theta peak central frequency and reduced its power without altering its bandwidth (Fig. 5 A-B). For the gamma peak, AlloP slowed the peak central frequency with no changes to power or bandwidth (Fig 5 A-B). AlloP also promoted emergence of a beta peak (Fig. 5A). Contrary to expectations for disinhibition, AlloP increased the aperiodic exponent and did not change the offset (Fig. 5 C-D). The lack of effect on offset is nominally distinct from pentobarbital and diazepam, though our studies, which focused on aperiodic exponent, were not powered to detect offset differences. The increase in exponent is inconsistent with the hypothesis that AlloP disinhibits cortical circuitry. However, the result is consistent with the effect of other GABA_A_R PAMs (Figs. 1, 2).

**Figure 5.**
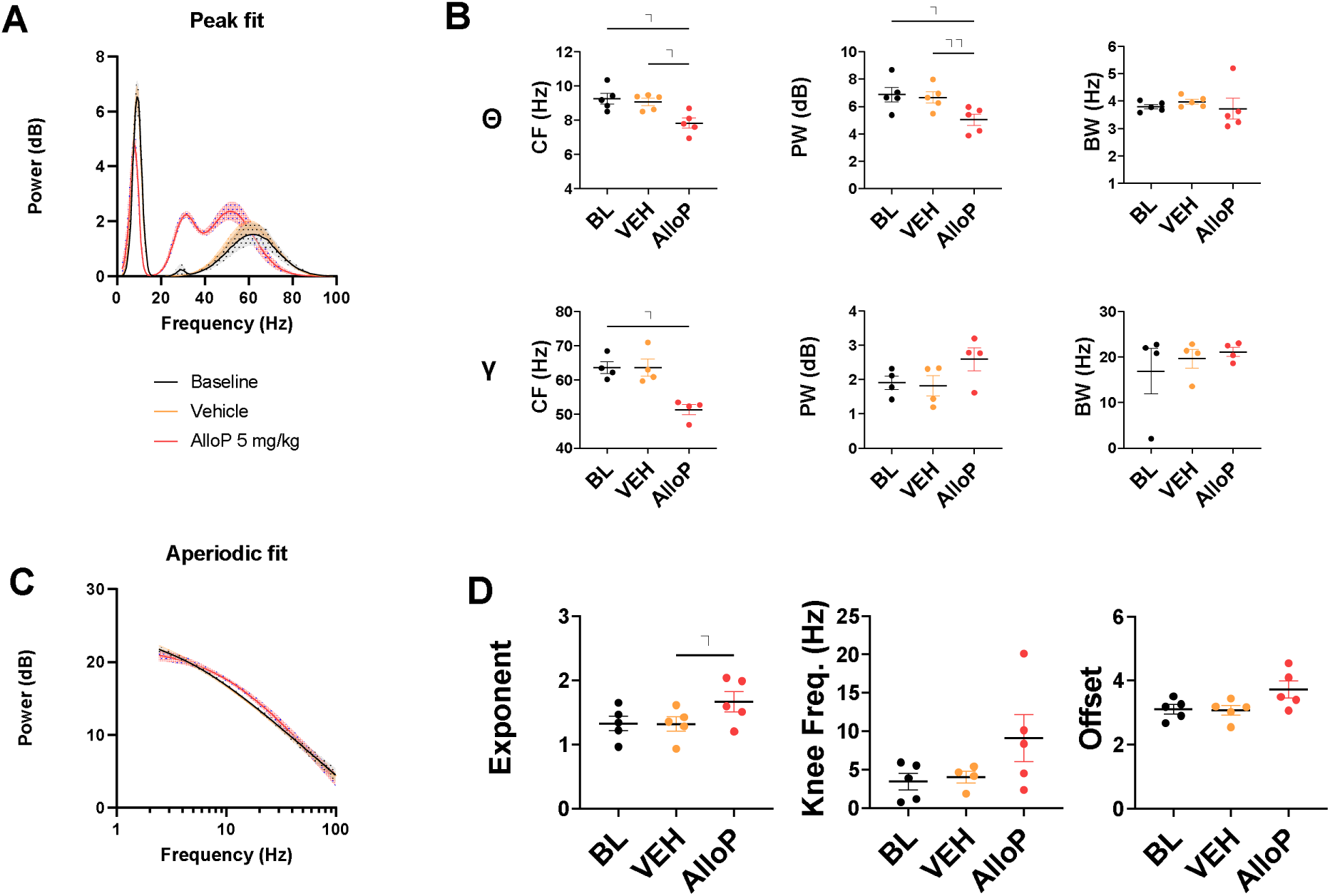
Allopreganolone 5 mg/kg increases aperiodic exponent, virtually indistinguishable from pentobarbital. **(A)** Periodic fit of peaks from baseline (BL), vehicle (VEH), and allopregnanolone (AlloP) effects on parietal active wake EEG raw power spectra. Emergence of a beta (12-30) peak with AlloP. **(B)** Parameters of theta (1-5) and gamma (30-100) peaks: peak central frequency (CF), peak power (PW) and peak bandwidth (BW). For the theta peak, a repeated measures one-way ANOVA revealed an effect of treatment on CF (F (1.471, 5.883) = 15.12 p = 0.0061). Tukey’s multiple comparison showed a difference between BL and AlloP CF (*p = 0.0266) and VEH and AlloP CF (*p = 0.0360). Repeated measures one-way ANOVA revealed an effect of treatment on PW (F (1.065, 4.260) = 27.57 p = 0.0051). Tukey’s multiple comparison showed a difference between BL and AlloP PW (*p = 0.0171) and VEH and AlloP PW (**p = 0.0055). Repeated measures one-way ANOVA revealed no effect of treatment on BW (F (1.059, 4.236) = 0.4042 p = 0.5692). For the gamma peak, a repeated measures one-way ANOVA revealed an effect of treatment on CF (F (F (1.631, 4.894) = 12.12 p = 0.0141). Tukey’s multiple comparison showed a difference between BL and AlloP CF (*p = 0.0214). Repeated measures one-way ANOVA revealed no effect of treatment on PW (F (1.377, 4.132) = 6.293 p = 0.0603). Repeated measures one-way ANOVA revealed no effect of treatment on BW (F (1.167, 3.500) = 0.3905 p = 0.6015). **(C)** Aperiodic fit of baseline, vehicle, and allopregnanolone effects on parietal active wake EEG raw power spectra. **(D)** Parameters of aperiodic fit: exponent, knee frequency (knee freq.), and offset. Repeated measures one-way ANOVA revealed an effect of treatment on exponent (F (1.064, 4.255) = 13.11 p = 0.0196). Tukey’s multiple comparison showed a difference between VEH and AlloP exponent (*p = 0.0408). Repeated measures one-way ANOVA revealed an effect of treatment on offset (F (1.082, 4.328) = 10.62 p = 0.0269). Tukey’s multiple comparison showed no differences between any treatment’s offset.

### Chemogenetic disinhibition fails to decrease aperiodic exponent

To shift cortical E/I ratio more definitively toward excitation and a hypothesized aperiodic exponent increase, we turned to cell-specific manipulations with DREADDs. We expressed hM4D(Gi) in parvalbumin-expressing (PV) interneurons, to dampen inhibition and thereby increase cortical activity upon administration of the designer ligand DCZ (Courtin et al., 2013; Yang et al., 2017; Yeganeh et al., 2022).Viral expression was widespread (Extended Data Fig. 6-1A). In cortices of the 3 mice used for expression analysis, we observed good overlap of mCherry viral expression with endogenous PV (Extended Data Fig. 6-1B), and 53.2 ± 5.3% (mean ± SEM) of PV cells were labeled with the mCherry viral reporter. hM4D(Gi) activation with DCZ produced strong effects on the power spectrum of parietal EEG (Extended Data Fig. 3-1E) with no discernible behavioral effect. Meanwhile, vehicle EEG in the same animals showed similar oscillatory and non-oscillatory components as baseline and vehicle controls in other experiments (Fig. 6, black symbols). DCZ slowed theta peak central frequency and reduced its power without changing bandwidth (Fig. 6 A-B). DCZ slowed the peak central gamma frequency with no changes to power or bandwidth (Fig. 6B). In contrast to expectations, chemogenetic suppression of PV interneurons with DCZ increased the exponent and offset of the aperiodic component of power spectra (Fig. 6 C-D). We also confirmed that DCZ did not alter the power spectra of control wild-type mice uninfected with virus (Extended Data Fig. 6-2).

**Figure 6.**
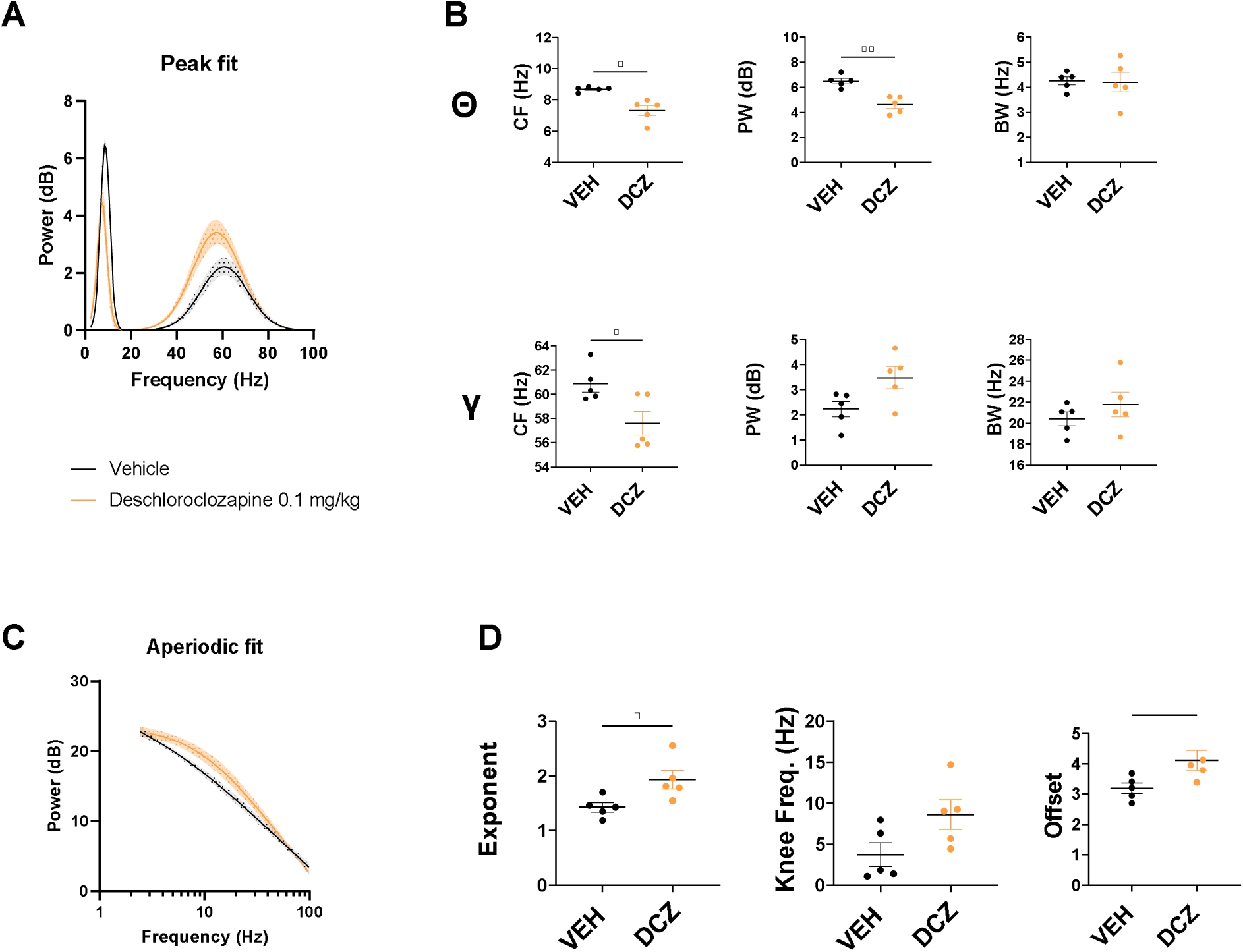
Chemogenetic inhibition of PV interneurons increases aperiodic exponent. **(A)** Periodic fit of peaks from baseline (BL), vehicle (VEH), and deschloroclozapine (DCZ) effects on parietal active wake EEG raw power spectra of hM4D(Gi) PV cre mice. **(B)** Parameters of theta (1-5) and gamma (30-100) peaks: peak central frequency (CF), peak power (PW) and peak bandwidth (BW). For the theta peak, a paired t-test, *t(*4) = 3.744 revealed a difference between VEH and DCZ CF (*p = 0.0200). A paired t-test, *t(*4) = 5.332 revealed a difference between VEH and DCZ PW (**p = 0.0060). A paired t-test revealed no effect of DCZ on theta BW. For the gamma peak, a paired t-test, *t(*4) = 3.887 revealed a difference between VEH and DCZ CF (*p = 0.0177). Paired t-tests revealed no effect of DCZ on gamma PW or BW. **(C)** Aperiodic fit of baseline, vehicle, and deschloroclozapine effects on parietal active wake EEG raw power spectra of hM4D(Gi) PV cre mice. **(D)** Parameters of aperiodic fit: exponent, knee frequency (knee freq.), and offset. A paired t-test, *t(*4) = 2.915 revealed a difference between VEH and DCZ exponent (*p = 0.0435). A paired t-test, *t(*4) = 3.235 revealed a difference between VEH and DCZ offset (*p = 0.0318).

We manipulated cortical PV cell activity in the opposite direction with hM3D(Gq) expression. We expressed hM3D(Gq) in PV interneurons with the same strategy used for PV suppression. Chemogenetic activation of PV cells with DCZ strongly suppressed EEG power across frequencies (Extended Data Fig. 3-1F) but also strongly arrested locomotion immediately upon injection, a state that typically lasted for ∼2-3 h (Extended Data Fig. 6-3A). The acute DCZ state interrupted ongoing oscillations across frequencies in a manner distinct from sleep states (Extended Data Fig. 3-1F vs. 3-1B; Extended Data Fig. 6-3B); thus oscillations during acute DCZ treatment could not be quantified (Extended Data Fig. 6-3 C-D). Upon recovery from sedation, oscillatory components were slightly altered relative to vehicle, possibly attributed to lower locomotion levels evident in the accelerometer (Extended Data Fig. 6-4A; C-D). Despite these oscillatory effects, the aperiodic components were unaffected relative to full wash out (Extended Data Fig. 6-3 E-F).

## Discussion

Here we explored the reliability of the aperiodic exponent of EEG power spectra as an indicator of E/I ratio. We compared the change in aperiodic exponent, extracted with FOOOF analysis, in various pharmacological manipulations in mice during active exploration. Overall, our results fail to uphold the hypothesis that the aperiodic exponent of EEG power spectra reliably marks E/I ratio. Along the way, we encountered other unexpected results.

To our knowledge, we are the first to report the unexpected sedation by GABA_A_ receptor anatgonists, picrotoxin and bicuculline, at sub-convulsive doses. Our results also highlight the importance of carefully considering behavioral state when comparing drug effects. Although some agents, notably GABA_A_R PAMs, produced changes to the aperiodic exponent in the expected direction (larger exponent signifying enhanced inhibition), other agents failed to change the aperiodic exponent in the predicted direction. In particular, sub-anesthetic ketamine failed to alter the aperiodic exponent despite robust effects on oscilliatory components. Although questions may be raised about the excitatory versus inhibitory impact of broadly acting drugs on cortical circuits, even selective chemogenetic disinhibition contradicted the hypothesis. Thus, we conclude that the aperiodic exponent is not a universally reliable measure of E/I ratio.

Picrotoxin and bicuculline’s sedative effects at sub-convulsive doses were surprising, as these GABA_A_R antagonists are classically thought to produce widespread excitation and are used to model seizure activity, a classic example of E/I increase (Fisher, 1989; Rohrbacher et al., 1998). However, an *in vitro* study demonstrated that picrotoxin reduces gamma oscillations in the rat primary motor cortex (Johnson et al., 2017), which may be necessary for active wake behavior.. This effect could be related to the paradoxical sedation and increased aperiodic exponent. Nevertheless, the unexpected results induced by GABA_A_R antagonism complicated efforts at EEG analysis and prompted us to query whether behavioral state itself is associated with changes to the aperiodic component of power spectra. Sedative-hypnotic effects of anesthetics share features with sleep. Although sleep and anesthesia are distinct states that have unique EEG patterns (Date et al., 2020), REM and NREM sleep both increased the aperiodic exponent (Extended Data Fig. 3-3), consistent with the effects of GABA_A_R PAMs and with the idea that sleep states also represent states of enhanced inhibition. However, because a classically excitatory drug (picrotoxin) also increased sedation and the EEG aperiodic exponent, we redoubled efforts to limit analysis to active wake. We first did this by examining the recovery from picrotoxin sedation (Fig. 3D), where the aperiodic exponent of picrotoxin during active wake did not change relative to baseline or vehicle, inconsistent with cortical excitation. Overall, caution may be warranted in interpreting the aperiodic exponent in studies that compare an anesthetized state with a resting wake state (Gao et al., 2017; Waschke et al., 2021), where confounding variables associated with the behvioral state difference may influence the aperiodic exponent.

Ketamine and hM3D(Gq) DREADD activation of PV interneurons caused no change in the aperiodic exponent when tested at sub-anesthetic doses or upon recovery from behavioral arrest. For ketamine, one interpretation might be that the disinhibition hypothesis of ketamine action is incorrect. However, given the robust effect of these treatments on oscillatory components, it seems unlikely that the treatments failed to produce either excitation or inhibition. Another issue is that disinhibition in rodents has mainly been demonstrated in prefrontal cortex or hippocampus rather than parietal cortex (Monyer et al., 1994; Moghaddam et al., 1997; Seamans, 2008; Zanos and Gould, 2018). Nevertheless, human data show ketamine-induced decreases in the aperiodic component in auditory and visual cortices, suggesting a widespread effect (Waschke et al., 2021). Thus, we suggest that ketamine and hM3D(Gq) DREADD activation of PV interneurons fail to support the primary hypothesis that the aperiodic exponent predicts E/I ratio.

AlloP, a GABA_A_R PAM and rapid acting antidepressant, disinhibits the baslolateral amygdala circuit by cell-selective actions (Antonoudiou et al., 2022). Here we found that AlloP increased rather than decreased the aperiodic exponent or cortical EEG, consistent with the other GABA_A_R PAMs (pentobarbital and diazepam). It is possible that cortical inhibition dominates the effect of AlloP during active wake while disinhibition is limited to basolateral amygdala. Alternatively, the aperiodic exponent may not be a good marker of AlloP induced disinhibition. It may be noteworthy that AlloP failed to increase aperiodic offset parameter, unlike pentobarbital and diazepam (Figure 1C, 2D, 5D), although AlloP caused trend-level increase. Because our hypothesis focused on the aperiodic exponent as a signature of E/I ratio, we did not power studies to explore the offset parameter. The idea that offset may provide a distinguishing feature among GABA PAMs remains to be rigorously tested.

Arguably our most definitive test of the primary hypothesis used cell-type selective supression of GABAergic interneurons, a well validated means of increasing activity in the cortex (Courtin et al., 2013; Yang et al., 2017; Yeganeh et al., 2022). DREADD mediated circuit disinhibition changed the aperiodic exponent in the direction opposite the hypothesis (Fig. 6D). A caveat is that PV positive cells throughout the brain were potentially affected, possibly complicating a purely cortical excitatory effect. Conversely, activation of PV interneurons (circuit inhibition), as expected, produced strong apparently sedative-hypnotic effects (Ferrari et al., 2022). The impact on EEG was so strong that poor FOOOF fits of sedative power spectra resulted, complicating efforts to interpret the impact of PV interneuron activation on the aperiodic exponent.

Overall, we failed to see a change in aperiodic exponent in the expected direction for excitatiory or disinhibitory manipulations including picrotoxin, ketamine, and DREADDs. Although GABA_A_R PAMs like pentobarbtal, diazepam, and AlloP all increased the exponent as expected, it remains unclear whether all forms of inhibition will be detected by the aperiodic exponent. Given the exceptions to the hypothesis, it is possible that E/I ratio as a construct and/or EEG-based measures of E/I ratio are too oversimplified to serve as an outcome prediction for a hypothesis. Regardless, our data do not support the aperiodic exponent of cortical EEG power spectra as a reliable marker of the concept of E/I ratio.

## Acknowledgements

The authors thank members of the Taylor Family Institute for Innovative Psychiatric Research for discussion and input. Widefield imaging was performed in the Washington University Center for Cellular Imaging (WUCCI).

## Author contributions

Conceptual study design: SVS, PML, & SM. Data collection and analysis: SVS, PL, AB, & NR. Original draft: SVS. Critical revisions: SVS, PML, NR, MW, CFZ, & SM.

