## Extended Data for "Periodic and aperiodic changes to cortical EEG in response to pharmacological manipulation"

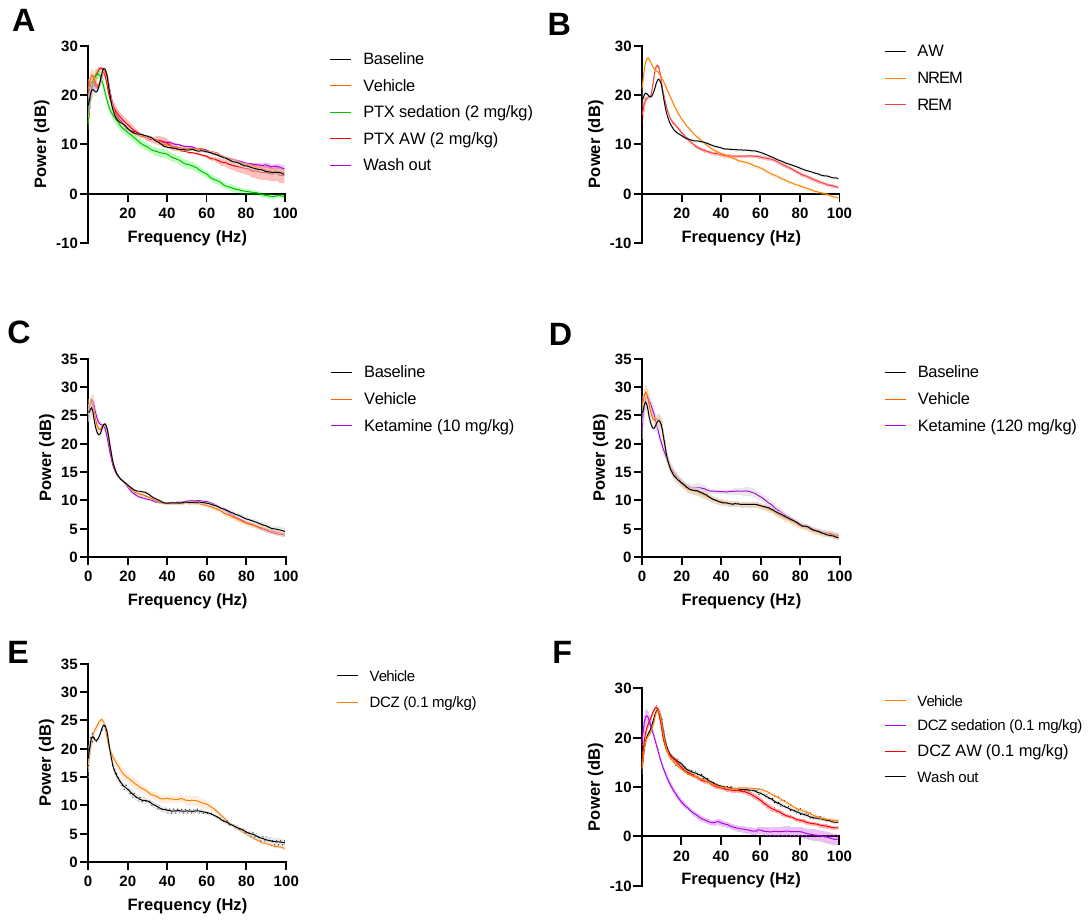

**Extended Data Figure 3-1. Raw power spectra of new treatment responses.**

Raw parietal EEG power spectra of treatment responses in EEG data presented in Figs. 3-4, 6, and Extended Data Figs. 2-4. (A- picrotoxin, B- behavioral states, C- sub-anesthetic ketamine, D- anesthetic ketamine, E- DCZ in PV hM4D(Gi) expressing mice, F- DCZ in PV hM3D(Gq) expressing mice).

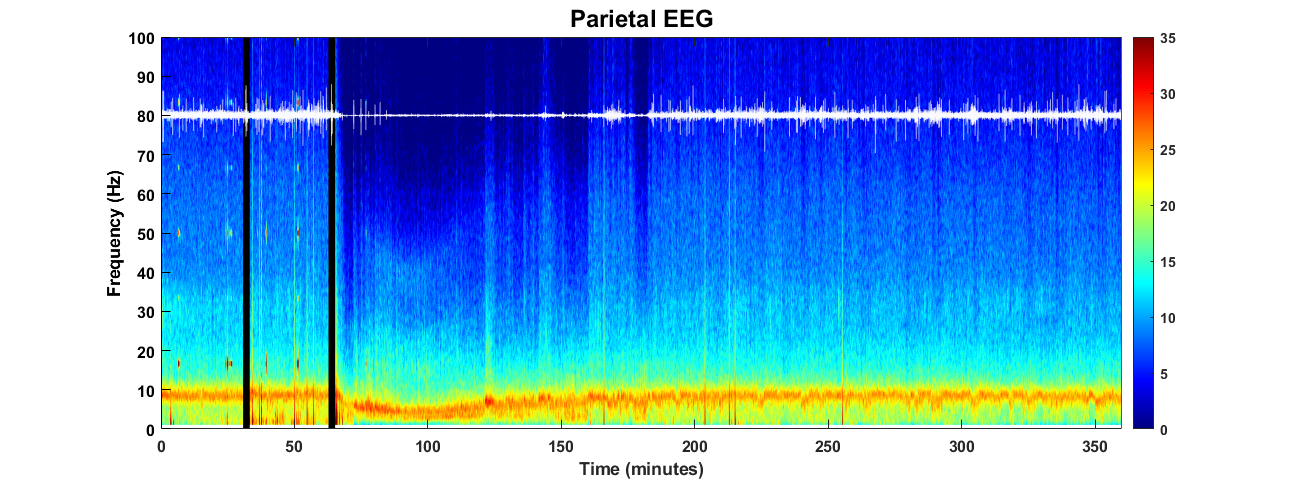
 **Extended Data Figure 3-2. Picrotoxin 2 mg/kg creates unexpected sedation.**

Representative spectrogram of picrotoxin’s sedative effects. Black vertical bars represent vehicle and picrotoxin injections (respectively). White overlay represents animal activity measured via accelerometer.

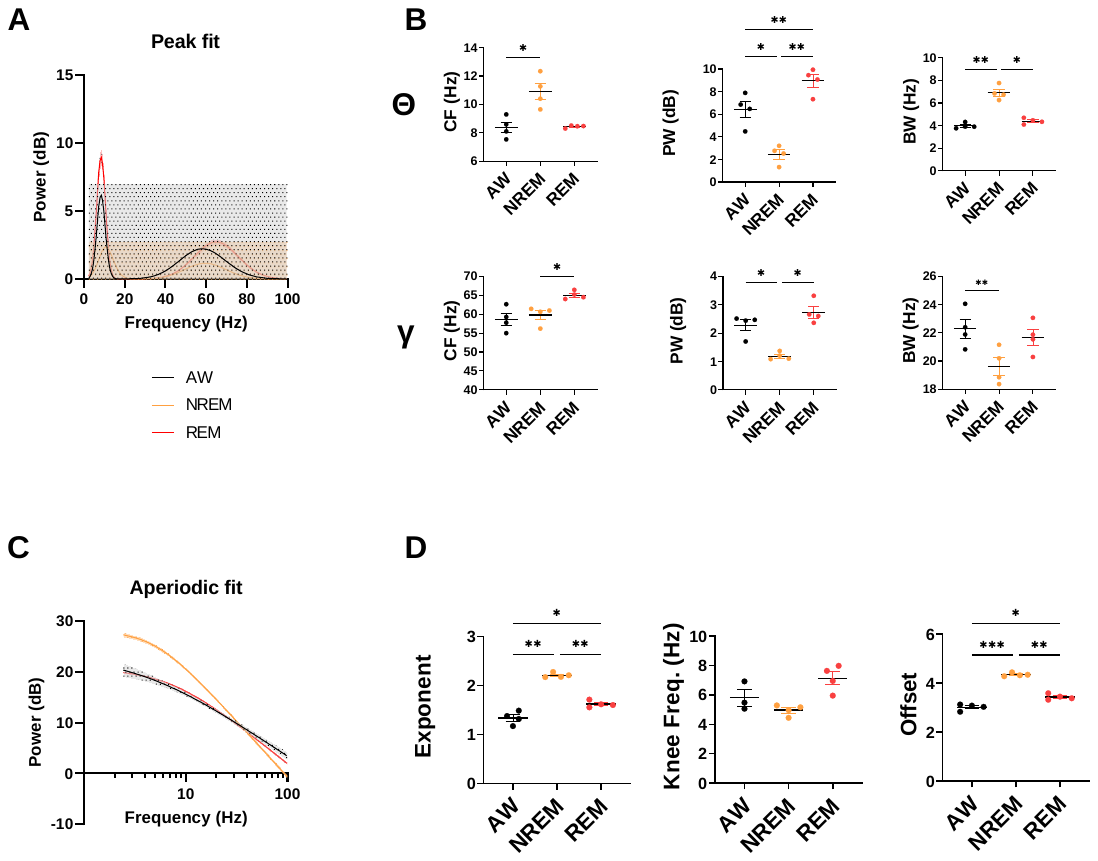
 **Extended Data Figure 3-3. Behavioral state changes exponent.**

**(A)** Periodic fit of peaks from active wake (AW), non-rapid eye movement (NREM), and rapid-eye movement (REM) effects on parietal active wake EEG raw power spectra. **(B)** Parameters of theta (1-5) and gamma (30-100) peaks: peak central frequency (CF), peak power (PW) and peak bandwidth (BW). For the theta peak, a repeated measures one-way ANOVA revealed an effect of behavioral state on CF (F (1.591, 4.772) = 15.04 p = 0.0098). Tukey’s multiple comparison showed a difference between AW and NREM CF (*p = 0.0452). Repeated measures one-way ANOVA revealed an effect of behavioral state on PW (F (1.026, 3.078) = 40.54 p = 0.0072). Tukey’s multiple comparison showed a difference between AW and NREM PW (*p = 0.0487), AW and REM PW (**p = 0.0014), and NREM and REM PW (**p = 0.0086). Repeated measures one-way ANOVA revealed an effect of behavioral state on BW (F (1.586, 4.758) = 49.59 p = 0.0008). Tukey’s multiple comparison showed a difference between AW and NREM BW (**p = 0.0076) and NREM and REM BW (*p = 0.0122). For the gamma peak, a repeated measures one-way ANOVA revealed an effect of behavioral state on CF (F (1.541, 4.622) = 15.10 p = 0.0107). Tukey’s multiple comparison showed a difference between NREM and REM CF (*p = 0.0268). Repeated measures one-way ANOVA revealed an effect of behavioral state on PW (F (1.685, 5.055) = 35.37 p = 0.0012). Tukey’s multiple comparison showed a difference between AW and NREM PW (*p = 0.0160) and NREM and REM PW (*p = 0.0130). Repeated measures one-way ANOVA revealed a trend toward effect of behavioral state on BW (F (1.025, 3.076) = 9.086 p = 0.0551). Tukey’s multiple comparison showed a difference between AW and NREM BW (**p = 0.0018). **(C)** Aperiodic fit of active wake, NREM, and REM effects on parietal active wake EEG raw power spectra. **(D)** Parameters of aperiodic fit: exponent, knee frequency (knee freq.), and offset. Repeated measures one-way ANOVA revealed an effect of behavioral state on exponent (F (1.579, 4.737) = 168.0 p < 0.0001). Tukey’s multiple comparison showed a difference between AW and NREM exponent (**p = 0.0014), AW and REM exponent (*p = 0.0214), and NREM and REM exponent (**0.0010). Repeated measures one-way ANOVA revealed an effect of behavioral state on offset (F (1.984, 5.951) = 151.9 p < 0.0001). Tukey’s multiple comparison showed a difference between AW and NREM offset (***p = 0.0008), AW and REM offset (*p = 0.0262), and NREM and REM offset (**p = 0.0031).

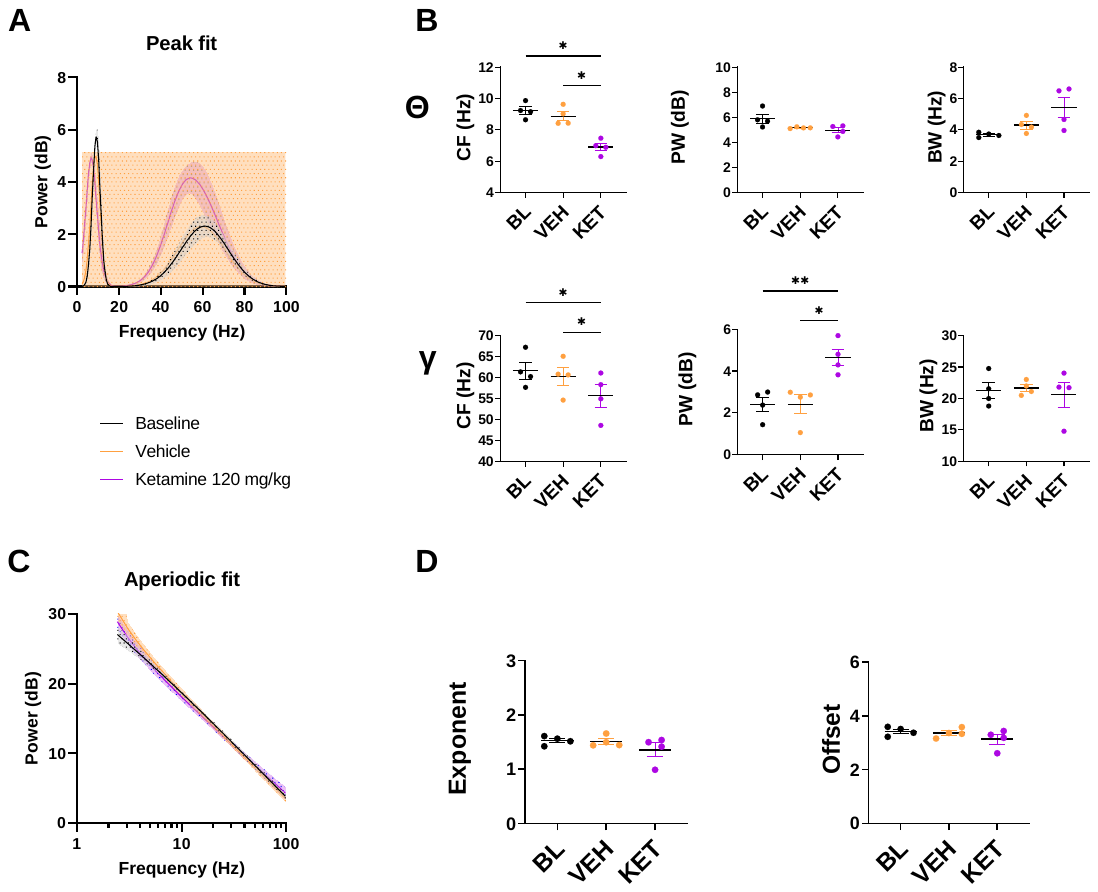

**Extended Data Figure 4-1. Anesthetic dose of ketamine (120 mg/kg) does not alter exponent.**

**(A)** Periodic fit of peaks from baseline (BL), vehicle (VEH), and ketamine (KET) effects on parietal active wake EEG raw power spectra. **(B)** Parameters of theta (1-5) and gamma (30-100) peaks: peak central frequency (CF), peak power (PW) and peak bandwidth (BW). For the theta peak, a repeated measures one-way ANOVA revealed an effect of treatment on CF (F (1.135, 3.406) = 28.80 p = 0.0087). Tukey’s multiple comparison showed a difference between BL and KET CF (*p = 0.0226) and VEH and KET CF (*p = 0.0270). Repeated measures one-way ANOVA revealed no effect of treatment on PW (F (1.014, 3.042) = 3.195 p = 0.1708). Repeated measures one-way ANOVA revealed no effect of treatment on BW (F (1.109, 3.327) = 3.794 p = 0.1381). For the gamma peak, a repeated measures one-way ANOVA revealed an effect of treatment on CF (F (1.409, 4.226) = 20.22 p = 0.0081). Tukey’s multiple comparison showed a difference between BL and KET CF (*p = 0.0362) and VEH and KET CF (*p = 0.0234). Repeated measures one-way ANOVA revealed an effect of treatment on PW (F (1.428, 4.285) = 58.29 p = 0.0009). Tukey’s multiple comparison showed a difference between BL and KET PW (**p = 0.0050) and VEH and KET PW (*p = 0.0101). Repeated measures one-way ANOVA revealed a trend towards no effect of treatment on BW (F (1.252, 3.755) = 0.1938 p = 0.7359). **(C)** Aperiodic fit of baseline, vehicle, and ketamine effects on parietal active wake EEG raw power spectra. **(D)** Parameters of aperiodic fit: exponent and offset. Repeated measures one-way ANOVA revealed no effect of treatment on exponent (F (1.187, 3.562) = 2.418 p = 0.2079). Repeated measures one-way ANOVA revealed no effect of treatment on offset (F (1.187, 3.562) = 2.418 p = 0.2079). No knee frequency reported, as all knees were negative but near 0.

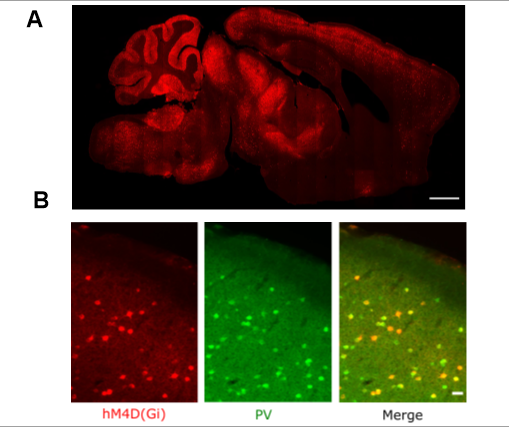

**Extended Data Figure 6-1. DREADD expression.**

**(A)** hM4D(Gi) expression in a sagittal brain slice. Scale bar 1000 μm. **(B)** Cortical expression of hM4D(Gi) (mCherry) is specific to PV interneurons (GFP). Scale bar 25 μm.

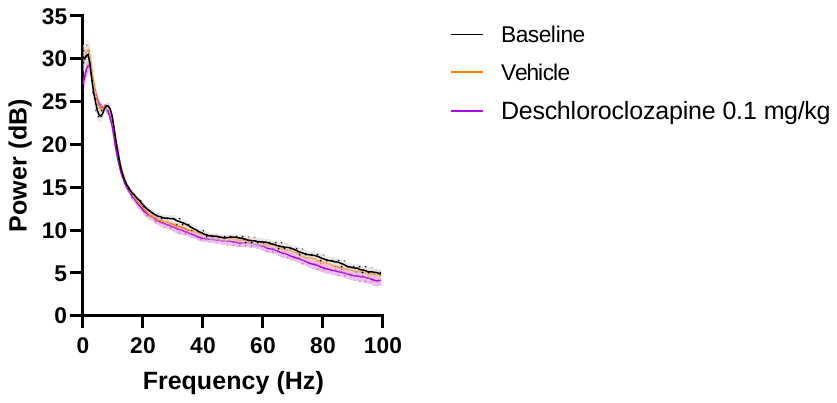

**Extended Data Figure 6-2. DCZ 0.1 mg/kg has no effect on wild type mice.**

Raw power spectra of baseline, vehicle, and deschloroclozapine (DCZ) 0.1 mg/kg in wild type mice reveals no effect of DCZ.

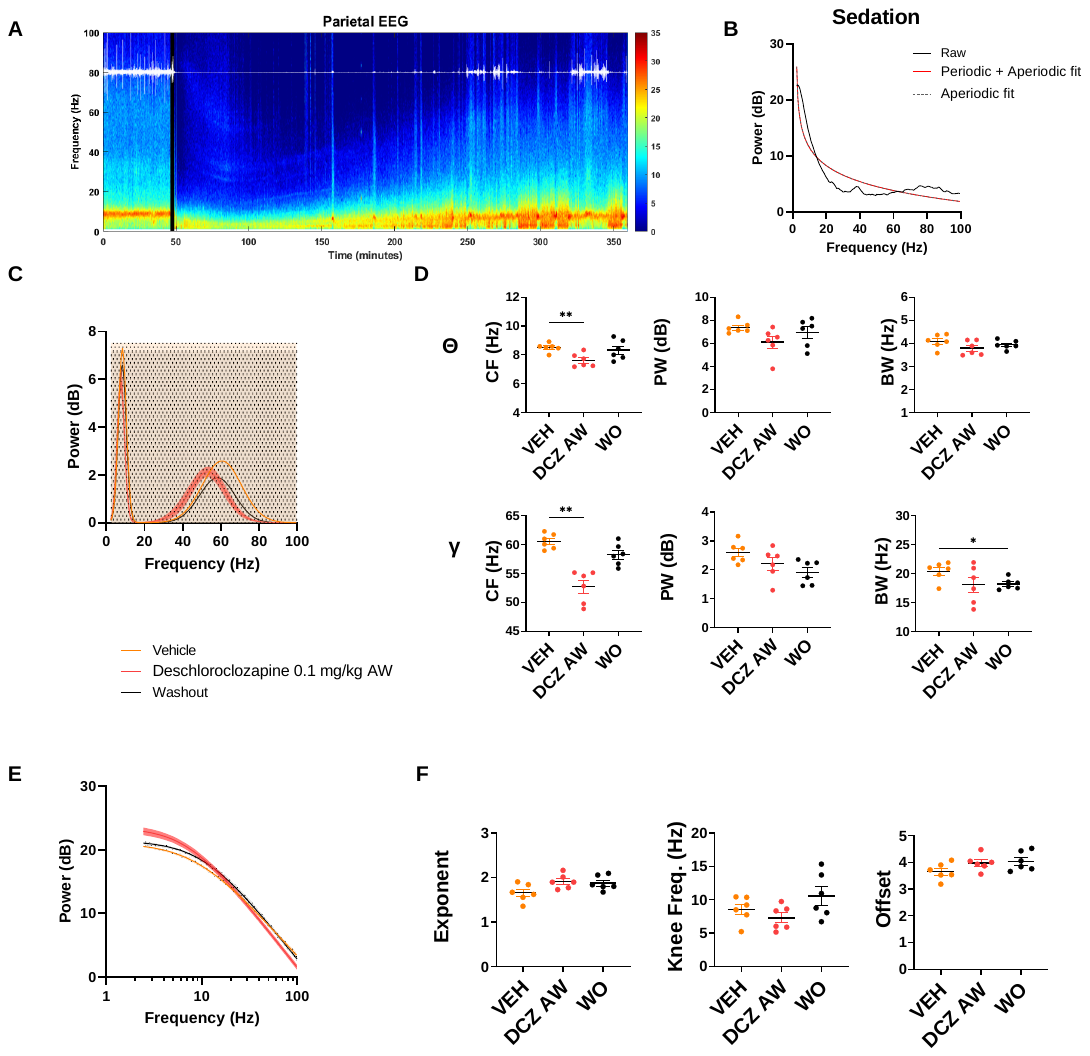
 **Extended Data Figure 6-3. Excitation of PV interneurons with hMD3(Gq) results in powerful sedation.**

**(A)**Representative spectrogram of DCZ’s sedative effects on hMD3(Gq) in PV interneurons. Black vertical bar represents DCZ injection. White overlay represents animal activity measured via accelerometer. **(B)** Representative DCZ sedation disrupted ongoing oscillations such that none could be accurately fit. **(C)** Periodic fit of peaks from baseline (BL), vehicle (VEH), deschloroclozapine (DCZ) active wake (AW), and washout (WO) effects on parietal active wake EEG raw power spectra of hM3D(Gq) in PV interneurons. **(D)** Parameters of theta (1-5) and gamma (30-100) peaks: peak central frequency (CF), peak power (PW) and peak bandwidth (BW). For the theta peak, a repeated measures one-way ANOVA revealed an effect of treatment on CF (F (1.275, 6.374) = 9.110 p = 0.0185). Tukey’s multiple comparison showed a difference between VEH and DCZ AW CF (**p = 0.0046). Repeated measures one-way ANOVA revealed no effect of treatment on PW F (1.181, 5.906) = 2.291 p = 0.1833). Repeated measures one-way ANOVA revealed no effect of treatment on BW (F (1.763, 8.816) = 3.530 p = 0.0786). For the gamma peak, a repeated measures one-way ANOVA revealed an effect of treatment on CF (F (1.235, 6.176) = 20.50	p = 0.0030). Tukey’s multiple comparison showed a difference between VEH and DCZ AW CF (**p = 0.0019). A repeated measures one-way ANOVA revealed an effect of treatment on PW (F (1.922, 9.612) = 5.099 p = 0.0319). Tukey’s multiple comparison showed no differences between treatments. A repeated measures one-way ANOVA revealed no effect of treatment on BW (F (1.188, 5.941) = 2.777 p = 0.1466). **(E)** Aperiodic fit of baseline, vehicle, deschloroclozapine active wake, and washout effects on parietal EEG raw power spectra of hM3D(Gq) in PV interneurons. **(E)** Parameters of aperiodic fit: exponent, knee frequency (knee freq.), and offset. Repeated measures one-way ANOVA revealed an effect of treatment on exponent (F (1.138, 5.689) = 6.680 p = 0.0413). Tukey’s multiple comparison showed no differences between treatments. Repeated measures one-way ANOVA revealed no effect of treatment on offset (F (1.538, 7.688) = 3.866 p = 0.0759).
